# Epg5 links proteotoxic stress due to defective autophagic clearance and epileptogenesis in *Drosophila* and Vici Syndrome patients

**DOI:** 10.1101/2024.04.23.590762

**Authors:** Celine Deneubourg, Hormos Salimi Dafsari, Simon Lowe, Aitana Martinez-Cotrina, David Mazaud, Seo Hyun Park, Virginia Vergani, Reza Maroofian, Luisa Averdunk, Ehsan Ghayoor-Karimiani, Sandeep Jayawant, Cyril Mignot, Boris Keren, Renate Peters, Arveen Kamath, Lauren Mattas, Sumit Verma, Arpana Silwal, Felix Distelmaier, Henry Houlden, Adam Antebi, James Jepson, Heinz Jungbluth, Manolis Fanto

## Abstract

Epilepsy is a common neurological condition that arises from dysfunctional neuronal circuit control due to either acquired or innate disorders. Autophagy is an essential neuronal housekeeping mechanism, which causes severe proteotoxic stress when impaired. Autophagy impairment has been associated to epileptogenesis through a variety of molecular mechanisms. Vici Syndrome (VS) is the paradigmatic congenital autophagy disorder in humans due to recessive variants in the ectopic P-granules autophagy tethering factor 5 (*EPG5*) gene that is crucial for autophagosome-lysosome fusion and ultimately for effective autophagic clearance. VS is characterized by a wide range of neurodevelopmental, neurodegenerative, and neurological features, including epilepsy. Here, we used *Drosophila melanogaster* to study the importance of *epg5* in development, ageing, and seizures. Our data indicate that proteotoxic stress due to impaired autophagic clearance and seizure-like behaviors correlate and are commonly regulated, suggesting that seizures occur as a direct consequence of proteotoxic stress and age-dependent neurodegenerative progression in *epg5 Drosophila* mutants, in the absence of evident neurodevelopmental abnormalities. We provide complementary evidence from *EPG5*-mutated patients demonstrating an epilepsy phenotype consistent with *Drosophila* predictions and propose autophagy stimulating diets as a feasible approach to control *EPG5*-related pharmacoresistant seizures.

## Introduction

Epilepsy is a common, debilitating and potentially lethal neurological disorder caused by increased neuronal network excitability that affects around 1% of the general population [1]. Epilepsy may be primarily genetic due to pathogenic variants in genes affecting neuronal excitability or acquired, for example secondary to trauma, infections or tumors. In addition, epilepsy is often found as a co-morbidity in a wide range of early-onset neurodevelopmental and adult-onset neurodegenerative diseases, including Alzheimer’s Disease (AD) [2].

Autophagy is an evolutionarily conserved lysosomal degradation pathway with fundamental roles in cellular homeostasis, infection defence and organelle quality control [3]. Probably reflective of its specific role in neuronal proteostasis, primary and secondary autophagy impairment has been demonstrated to occur in numerous neurodegenerative diseases, including AD, Amyotrophic Lateral Sclerosis(ALS)/Frontotemoporal Dementia (FTD) and Parkinson’s Disease (PD) [4,5]. In addition, pathogenic variants in genes directly affecting autophagy have been recently identified in a growing list of neurodevelopmental disorders with superimposed neurodegeneration later in life, collectively termed congenital disorders of autophagy [6,7].

Postmitotic neurons, with their vast axonal and dendritic projections, high energy metabolism and virtual absence of cell renewal by proliferation are especially dependent on autophagy for self-maintenance and survival, providing an explanation for the disproportionate impact of autophagy dysfunction on the nervous system in human disease.

The role of autophagy in epileptogenesis has received particular attention, starting from initial observations in *TSC*-related tuberous sclerosis (TS) resulting in aberrant regulation of the mammalian Target Of Rapamycin (mTOR) pathway and, consequently, both disturbed autophagy and seizures. Whilst there is thus a large body of evidence suggesting that abnormal autophagy is linked to epilepsy [8–13], most work to date has focussed on upstream autophagy regulators such as mTOR but to a much lesser extent on defects located further downstream in the autophagy cycle as a cause of neuronal pathology and epileptogenesis.

The paradigmatic disorder of congenital disorders of autophagy is *EPG5*-related Vici Syndrome (VS) [14–16], a severe multisystem disorder and the first condition in which a primary autophagy defect was identified as the underlying genetic cause. Recessive variants in the ectopic P-granules autophagy tethering factor 5 gene (*EPG5*), which encodes a tethering factor required for effective autophagosome-lysosome fusion in the final step of the autophagy pathway, were linked to the disorder in 2013 [17,18]. VS comprises a wide range of neurodevelopmental, neurodegenerative and neurological features, with early-onset epilepsies reported in around two thirds of patients [15].

Following the initial description of VS in 1988 and its genetic resolution in 2013, a rapidly increasing number of conditions have emerged within the novel class of congenital disorders of autophagy [7]. Despite considerable clinical variability, these disorders are typically characterized by a neurodegenerative component of variable onset superimposed on an early-onset neurodevelopmental disorder. The natural history of neurological features in VS and related congenital disorders of autophagy such as beta-propeller protein-associated neurodegeneration (BPAN) [19–22] raises thus the question if the epilepsy commonly observed in these conditions is simply secondary to the principal neurodevelopmental defect and/or driven by the superimposed neurodegenerative component.

We have previously used *Drosophila melanogaster* to investigate the function of the VS gene *epg5* (CG14299), showing that its knock down is sufficient to generate autophagy clearance defects during larval development and premature neuronal degeneration during ageing [15].

We have now used the fruit fly for in-depth investigation of the epileptic component associated with *EPG5* mutations in humans, based on the rationale that *Drosophila* is already used as an epilepsy model [23,24] and offers a valid and ethical replacement for severe procedures in vertebrate animals. Seizures in hyper-excitable fly mutants are robustly induced by mechanical, electric or thermic stimuli [25,26]. Based on novel methodologies for the measurement of seizure-like behaviour and neuronal activation in live flies, we show that mutations in *epg5* enhance sensitivity to seizure-inducing stimuli. In addition, we demonstrate that this effect is age-related and regulated by factors that modulate proteostatic clearance. Our data support a direct association between autophagy impairment, proteotoxic stress and epilepsy, which can arise in the absence of overt neurodevelopmental abnormalities. Our analysis in *Drosophila* also suggests autophagy-stimulating diets as potential non-pharmacological treatments to ameliorate the proteotoxic stress due to autophagic clearance impairment and achieve seizure control in patients with VS and related disorders. Finally, we also provide complementary clinical and neurophysiological evidence from a subset of 12 *EPG5*-mutated patients from 11 families, suggesting evolution of epilepsy later in life corresponding to disease progression, as well as preliminary data indicating responsiveness to specific treatment modalities including autophagy-stimulating diets.

## Results

### *epg5* loss-of-function reduces lifespan and causes age-dependent motor decline

Our previous work into the role of *epg5* in *Drosophila* concentrated on its effect on larval fat bodies and on adult photoreceptor neurons [15]. We have now used several UAS-inducible RNAi lines to address the full physiological effect of *epg5* knock down in the nervous system using the pan-neuronal *elav-Gal4* driver. Surprisingly, none of the tested RNAi lines induced overt neurodevelopmental deficits in the larvae. Similarly to the corresponding larvae, adult *epg5* knockdown flies eclosed from the pupal case at the expected Mendelian rate and appeared morphologically normal. However, upon ageing it became evident that expression of two out of three RNAi lines shortened lifespan (Fig S1A). In addition, all RNAi lines caused significant progressive decline in a motor climbing assay, in comparison to control flies (Fig S1B).

To address whether the lack of a developmental phenotype was due to the delay in triggering *epg5* knock down due to the Gal4/UAS system of expression and possible protein perdurance, we generated a full *epg5* knock out using CRISPR/Cas9 gene-editing, replacing the whole *epg5* gene with an *mCherry* knock-in construct (Fig 1A). Mutant flies were isogenized for 6 generations in *w^1118^* background and validated for the insertion by PCR (Fig S1C) and mCherry expression (Fig S2A).

**Figure 1.**
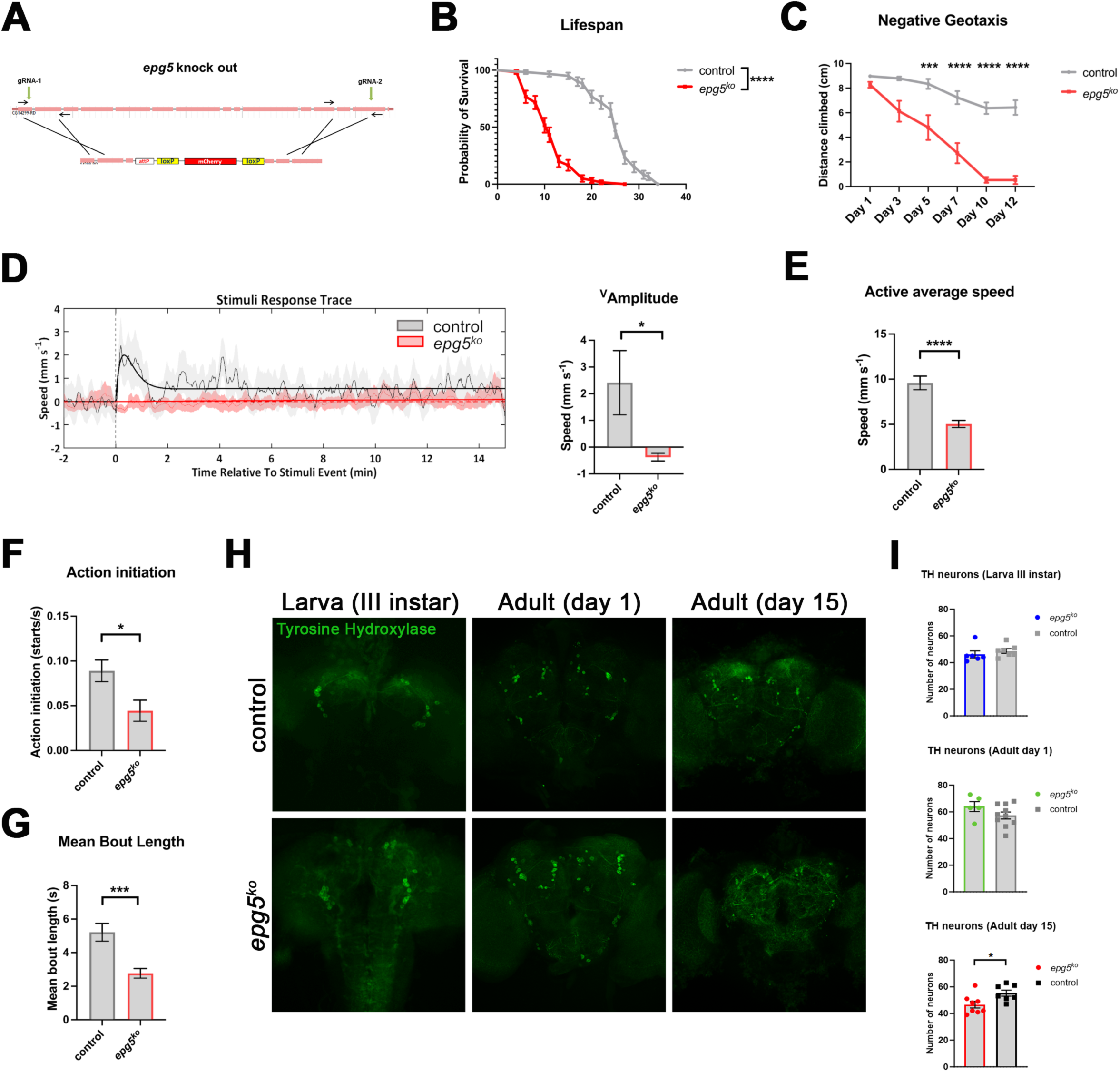
Age-related degeneration in *epg5* knock out flies. (A) Schematic representation of the *epg5* knock out generation. (B) Survival rate (%) of *epg5^ko^* flies in comparison to an isogenic control. N=60 flies per genotype. Kaplan Meier Log-rank test, ****p<0.0001. (C) Climbing assay of *epg5^ko^* and isogenic control flies. N=∼30 flies per genotype. Data are presented as mean ± S.E.M, Two-way ANOVA with Bonferroni’s multiple comparisons test. ***p<0.001, ****p<0.0001. (D-G) DART analysis of 7 days old *epg5^ko^* male flies and isogenic controls. Display reduced startling response to a train of vibrations (D), reduced speed when active (E), reduced number of actions initiated (F) and reduced length of each action bout (G). (H) Max intensity z-projections of confocal stacks through the brains of *epg5^ko^* and isogenic controls at different life stages stained for Tyrosine Hydroxylase to reveal dopaminergic neurons and quantification of the number of TH neurons per brain (I). Data are presented as mean ± S.E.M and were analysed using an unpaired Student’s t-test. *p<0.05, **p<0.01, ***p<0.001, ****p<0.0001.

In agreement with the observations with RNAi-mediated knock down, *epg5^ko^* flies did not display any overt neurodevelopmental abnormalities as third instar larvae and newly eclosed adults but had a shorter adult lifespan (Fig 1B). A significantly shorter lifespan was detected also when the *epg5^ko^* allele was placed in trans-heterozygosis over a deficiency covering the *epg5* locus (Fig. S1D). In addition, the knock out flies displayed a progressive decline in motor climbing ability in comparison to isogenic controls (Fig 1C). Interestingly, also heterozygous flies displayed similar lifespan and climbing defects, albeit at a lower level (Fig S1E,F) that was similar to the knock down effects.

When aged for 16 days at 29°C, *epg5^ko^* flies also displayed an increased accumulation of Ref(2)P, the ortholog of mammalian p62, puncta (Fig S2A,B) and of Atg8a puncta (Fig S2C,D), consistent with the expected defects in autophagic clearance.

To perform a more in-depth analysis of motor behaviour in 7 days old flies, the unbiased and automated *Drosophila* Arousal Tracking (DART) system was used. The DART assay allowed us to investigate the baseline activity of the flies, as well as their horizontal movements, and their response to an automated mechanical stimulus (vibration) [27–29]. This analysis focussing on aged male flies confirmed a defect in responding to stimuli with an increase in activity (Fig 1D) and indicated that *epg5^ko^* flies were significantly slower when active (Fig 1E), initiated fewer actions (Fig 1F) and had a much shorter action bout length (Fig 1G), suggesting a wide-ranging deficit in motor activity.

To analyse neurodevelopment and neurodegeneration in *epg5^ko^* flies at cellular level in specific neuronal populations, we have chosen two well defined neuronal circuitries within the fly brain. We have documented the evolution of dopaminergic neurons and mushroom body neurons from larval stages to early adulthood and endstage. To test for neurodevelopmental anomalies, we examined the structure of the adult mushroom bodies (MBs), a key learning and memory centre in Drosophila. We observed no clear alteration in the growth of axonal projections comprising the α and β lobes, nor any defects in axonal pathfinding, as evidenced by the lack of mid-line crossing of beta lobe axons (Fig S2E,F). Similarly we detected no significant alteration in the number of dopaminergic neurons in larval or early adult stages (Fig 1H,I). However, in ageing, whereas the α and β lobes of fibers in mushroom body neurons appeared unchanged (Fig S2E,F), the dopaminergic neurons number declined with age (Fig 1H,I), indicating neurodegeneration in *epg5* mutants.

In conclusion, modulating levels of *epg5* in *Drosophila* causes neurodegenerative features compatible with those seen in patients, while neurodevelopmental abnormalities are not detectable.

### *epg5* knock out sensitizes to seizure induction and neuronal hyperactivity

Given the high prevalence of seizures in VS patients we investigated whether *epg5^ko^* flies recapitulated this characteristic and how it evolved with ageing. In a validated hyperthermia-induced seizure assay [29,30] no relevant difference was observed between control and *epg5^ko^* flies one day post-eclosion (Fig 2A). When aged for 16 days at 29°C however, *epg5^ko^* flies developed a strong seizure-like behaviour induced by hyperthermia in comparison to control flies (Fig 2B). The flies that did show seizure-like behaviour also had a strongly increased recovery time compared to the control flies, specifically when aged (Fig 2C, and Fig S3A). The same effects were detected when the *epg5^ko^* allele was placed in trans-heterozygosis over a deficiency covering the *epg5* locus (Fig. S3B,C).

**Figure 2.**
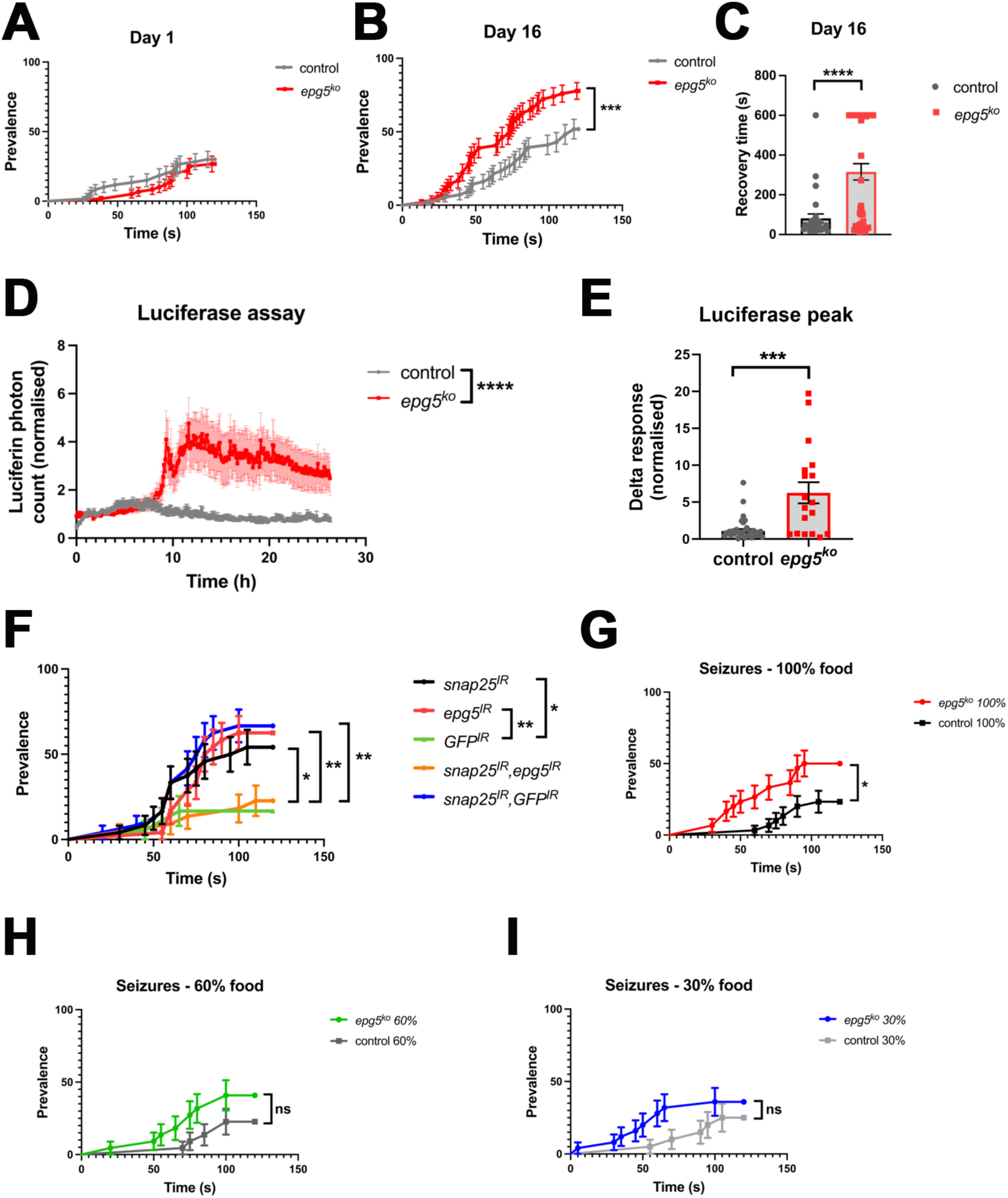
Age-related seizure-like behaviour in *epg5* knock out flies. (A-B) Seizure prevalence at day 1 (A) and day 16 (B) of age in *epg5^ko^* flies in comparison to an isogenic control. N=54-60 flies per genotype. Data are presented as mean ± S.E.M and analysed using the Log-rank (Mantel-Cox) test, ***p<0.001. (C) Recovery time of flies with seizures at day 16. N=29-42 flies per genotype. Data are presented as mean ± S.E.M and analysed using an unpaired Student’s t-test ****p<0.0001. (D) Average number of luciferin photons counted per genotype over 24 hours in the *in vivo* reporter system. N=18-36 flies per genotype. Data are presented as mean ± S.E.M and analysed using a two-way ANOVA with Bonferroni’s multiple comparisons test, ****p<0.0001. (E) The maximum number of photons counted per fly during the recording. N=18-36 flies per genotype. (F) Seizure prevalence of *snap25* and or *epg5* knock down in the nervous system and related controls. Data are presented as mean ± S.E.M. and are analysed using the Log-rank (Mantel-Cox) test. N=18-24 per genotype. *p<0.05, **p<0.01. (G-I) Seizure prevalence of 16 days old *epg5^ko^* flies in comparison to an isogenic control under normal food (G) and two caloric restriction conditions, 60% food (H) and 30% food (I) Data are presented as mean ± S.E.M and analysed using a Mann-Whitney test for, ***p<0.001.

Furthermore, to determine whether this effect was at least partly due to adult neurons in aged flies, we tested flies with RNAi-mediated knock down of *epg5* only in neurons after normal development. To achieve an adult specific knock down we used a Gal80^ts^ transgene (a temperature-sensitive inhibitor of the Gal4 transactivator) and transferred flies to a restrictive temperature of 29°C after eclosion to trigger *epg5* knock-down only from that stage. Resulting flies were again tested in the hyperthermia-induced seizure assay and showed an increased prevalence of seizures compared to control flies when aged for 29 days (Fig S3D). The recovery time was not significantly different from control flies, despite an obvious upwards trend (Fig S3E).

To examine if the seizure-like behaviour observed in flies is correlated with neuronal hyperactivity, we deployed an *in vivo* reporter system previously used to measure the activity of adult fly clock neurons [31,32]. This system utilises a fusion protein called CaLexA that consists of NFAT (nuclear factor of activated T-cells), which undergoes nuclear translocation in a Ca^2+^-dependent manner, and the bacterial transcription factor LexA [33]. Increases in intracellular Ca^2+^ drive nuclear localisation of CaLexA and enhance expression of a *luciferase* (*luc*) reporter gene downstream of the LexA binding site, LexAop (Fig. S3F). We expressed CaLexA in neurons of control and *epg5^ko^* flies fed with the Luciferase substrate D-Luciferin, subjected them to seizure-inducing hyperthermia, and quantified subsequent photon emission via luminometry (Fig S3G). Approximately 10 hours following hyperthermia - a time frame compatible with the known reaction time of the CaLexA-Luciferase system [31] – we observed a strong increase in photon count in the *epg5^ko^* flies compared to control flies, indicating elevated neuronal Ca^2+^ levels following seizure induction (Fig 2D). This can also be observed in the peak response compared to baseline response of *epg5^ko^* flies which is significantly higher than for control flies (Fig 2D,E).

Overall, these data indicate that both global and neuron-specific reduction of *epg5* expression sensitize *Drosophila* to seizure-inducing stimuli and that global loss of *epg5* evokes generalized neuronal hyperactivation.

### Autophagic clearance of proteotoxic stress and its regulation by *epg5* is directly linked to seizure-like behaviour

The function of Epg5 is that of a tethering factor facilitating specific fusion events between autophagosomes and lysosomes leading to autophagic clearance through interactions with specific SNARE complexes. In absence of Epg5, it has been shown that in mammalian cells abnormal STX17-SNAP25-VAMP8 SNARE complexes are formed. Interestingly, knock down of *SNAP25* together with *EPG5* ameliorated the autophagy clearance defects that is characteristic in *EPG5* deficiencies [18], suggesting that EPG5 deficiency only exerts its full pathogenic effect in the presence of a functional SNAP25 protein. In the context of epilepsy, *SNAP25* is highly relevant as its role in epileptogenesis is well-established and it has already been shown to be mutated in patients with a form of hereditary epilepsy [34,35]. We therefore studied the interaction between *snap25* and *epg5* in *Drosophila* using adult specific neuronal knock down. Interestingly, *snap25* knock down, but not a control GFP RNAi, elicited a robust seizure-like behaviour just like *epg5* knock down in flies aged 29 days (Fig 2F). Surprisingly, however, when combined, the concomitant knock-down of *epg5* and *snap25* drastically reduced the seizure prevalence (Fig 2F). A similar reduction of seizure prevalence does not occur when a control RNAi for GFP is used together with *snap25* knock down, indicating that this observation does not arise from the titration of the Gal4 transcription factor by an additional UAS promoter.

Importantly, the combination of *epg5* and *snap25* knock-down also reduced the proteotoxic stress, as detected by Ref(2)P accumulation (Fig. S3H). Therefore, the interaction between *epg5* and *snap25* in the regulation of seizures is similar to the interaction observed in autophagic clearance, implying a common basis for the sensitivity to seizures and autophagy in *epg5* mutants.

In consideration of the possible direct link between autophagy defects and seizures in *epg5^ko^* flies we tested whether caloric restriction could ameliorate the sensitivity to increased seizure via an increase in autophagic flux, considering the recognized role of fasting as a potent autophagy inducer. We produced a series of dilution of our standard fly-food recipe (60%, 30%, 10%, while 100% is full fly recipe), maintained newly eclosed adult flies for 9 days on this diet and then analysed for the accumulation of Ref(2)P to detect potential effects on autophagy in the brain. 10% food was insufficient to allow fly survival, including control flies, and was therefore abandoned. As shown previously, *epg5^ko^* flies fed on a standard diet (100% food) have significant accumulation of Ref(2)P particles compared to control flies fed the same fly food (Fig S3I). In flies that are mildly calorically restricted to 60% or 30% food the trend in the difference in the number of Ref(2)P particles that can be observed between *epg5^ko^* and control flies is reduced or no longer statistically significant (Fig S3I). Changing the nutrition of the flies by restricting their caloric intake thus seems to attenuate the defective autophagic clearance in *epg5^ko^* flies.

Interestingly, attenuation of autophagic clearance through mild fasting corresponded to a decrease in the sensitivity to hyperthermia-induced seizures. In particular, consistent with the Ref(2)P accumulation, feeding the flies 60% or 30% of the caloric value of standard fly food attenuated the increased prevalence of seizures of *epg5^ko^* flies, leading to a loss of statistical significance when compared to controls (Fig 2G-I).

A similar effect was also detected with regards to motor behaviour, where 30% food reduced the difference in climbing ability between *epg5^ko^* and control flies (Fig S3J).

Thus, caloric restriction ameliorates autophagic clearance defects caused by loss of *epg5* and partially rescues the seizure-like behaviour in *Drosophila*, arguing for this to be a direct consequence of the autophagy defect.

### *epg5* seizure-like behaviour can be detected with increased power though an automated DART-based platform

To standardise seizure behaviour analysis and make it unbiased and less dependent on investigator dexterity, we have developed a prototype machine that uses the DART software for seizure analysis. We have placed flies vertically in single tubes and fitted the tubes into an hydraulic chamber with fast water flow for induction of hyperthermia and video recording (Fig. S4A,B). Seizures are determined by a set of parameters regarding fly position and speed, with an arbitrary cut off for duration of events (see methods). In a proof-of-concept experiment, we have analysed 80 flies of both sexes for *epg5^ko^* and controls and established that the DART software detects a significant increase in the seizure-like behaviour of *epg5^ko^* flies with respect to controls (Fig 3A). In addition, the DART software is able to detect all seizure events identified in a single fly. This analysis indicates that a greater number of *epg5^ko^* flies experienced two or more seizing events when compared to controls (Fig 3B), while duration and strength of convulsions as detected by speped of movement without displacement were similar (Fig S4C). Therefore, when considering the seizing events in the populations analysed the *epg5^ko^* population displayed a much more significant prevalence of seizures compared to the control population (Fig, 3C).

**Figure 3.**
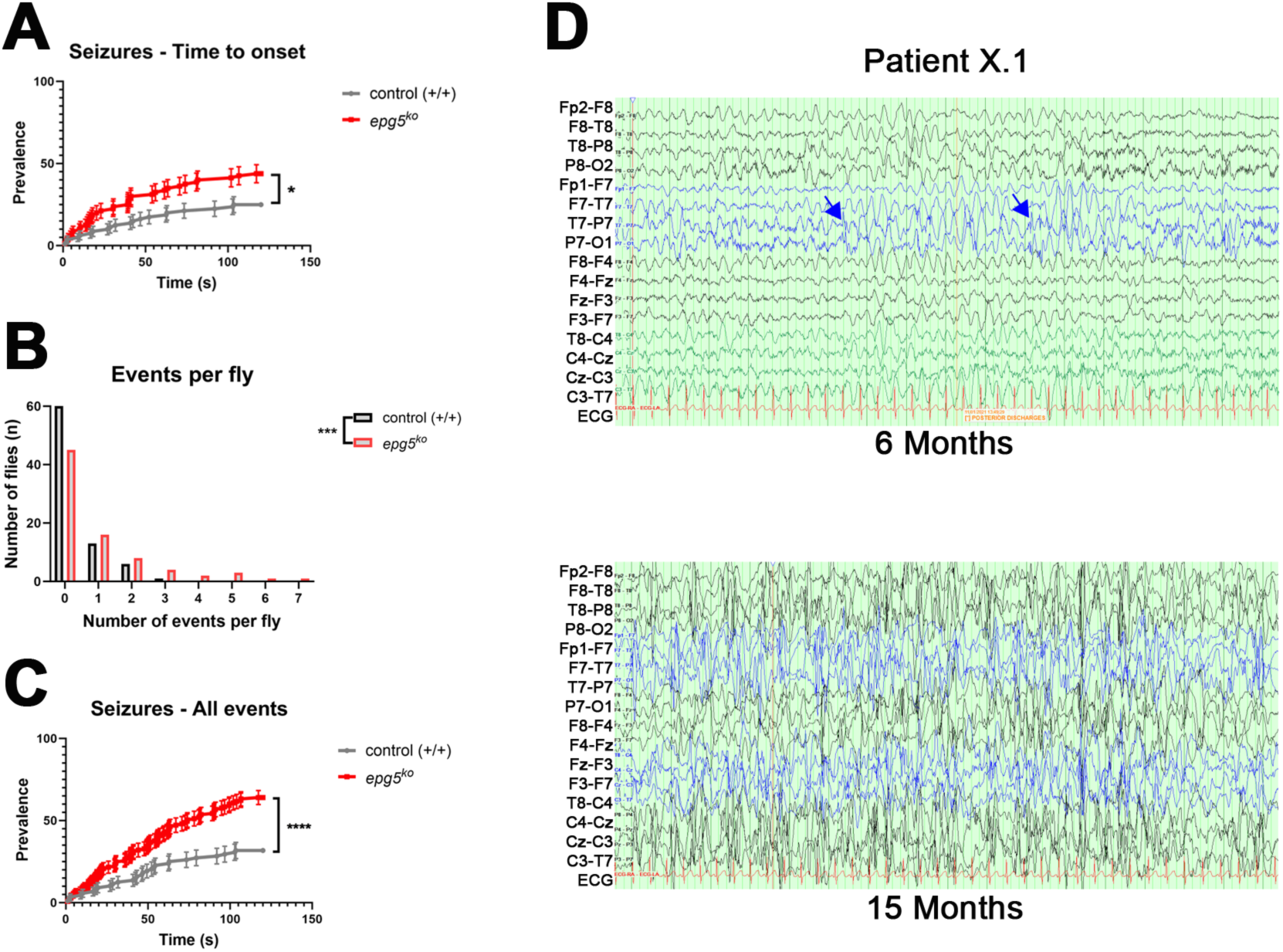
Analysis of seizures in *Drosophila* and in patients with *EPG5*-related epilepsy. (A) Automated epi-DART analysis of the time to onset of seizures (first seizure only). The *epg5^ko^* flies showed higher prevalence of seizing events compared to control flies. Data are presented as mean ± S.E.M and analysed using the Log-rank (Mantel-Cox) test *p<0.05, N=80 flies per group). (B) Number of seizing events per fly. Data are presented as contingency table and analysed through a Chi-square test for trend *χ^2^*= (1, *N*=80) =10.86, ***p<0.001, (C) Overall time of onset of all seizing events as detected by the automated DART analysis. Data are presented as mean ± S.E.M and analysed using the Log-rank (Mantel-Cox) test ****p<0.0001, N=88-125 events per group. (D) EEG recordings from Patient X.1 with posterior discharges maximum left parietooccipital and focal slowing at age of six months (top panel) and clear progression into hypsarrhythmia 9 months later (bottom panel).

Therefore, the seizure-like behaviour of *epg5^ko^* flies in can be detected automatically with our novel apparatus and the functions developed within DART with increased power for the discrimination of two genotypes.

### The epilepsy phenotype in several *EPG5*-mutated patients is of later onset and evolves over time

Whilst it has been recognized that a large proportion of VS patients have seizures [15], the *EPG5*- associated epilepsy phenotype has not yet been described in detail and its precise aetiology is currently not well-understood.

Epilepsy seen in *EPG5*-mutated patients may be a reflection of the underlying neurodevelopmental defect, and/or, as suggested by our *Drosophila* models, secondary to superimposed neurodegenerative changes that develop over time.

Prompted by our observations in *Drosophila* and to further delineate the *EPG5*-associated epilepsy phenotype in humans, we reviewed in detail the epilepsy phenotype and electroencephalogram (EEG) findings in 12 patients with pathogenic variants in *EPG5* (Table 1) identified through an international collaboration dedicated to *EPG5*-related disorders. These patients had onset of epileptic seizures between late infancy and pre-school age, except two cases where seizures (probably due to hypoxic-ischemic encephalopathy in at least one) were already noted in the neonatal period but completely ceased until onset of distinct epileptic phenomena later in childhood. One patient presented with typical infantile spasms from 15 months of age. Non-regarding the seizure phenotype at onset, in most patients, seizures changed into generalized spasms, tonic or atonic seizures over time, often exacerbated during infections.

**Table 1.**
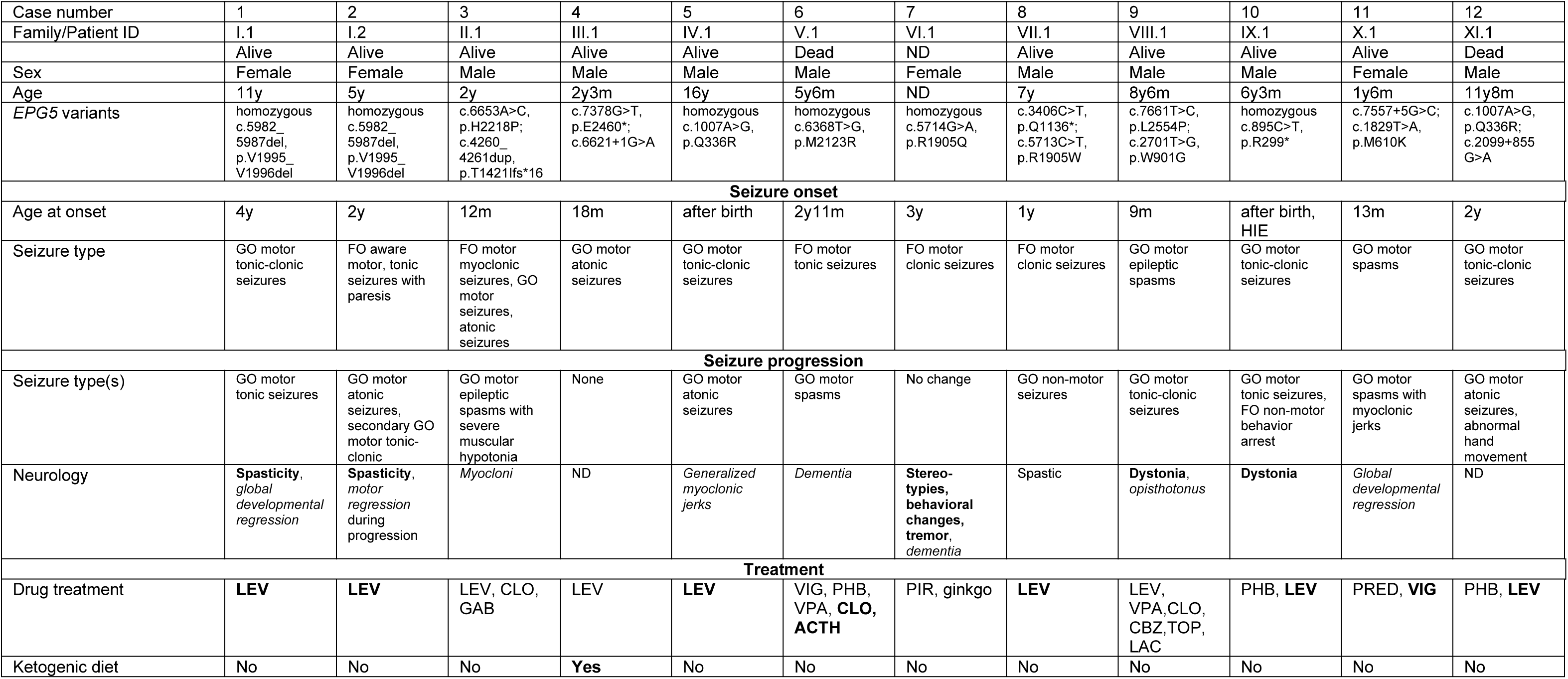
Characterization of seizure phenotypes, corresponding EEG abnormalities and treatments in patients with *EPG5*-related epilepsy. Age indicates age at last follow-up exam (if alive) or age of death. Genetic and protein alterations are indicated according to ENST00000282041.11, NM_020964.3 and ENSP00000282041.4, NP_066015.2), respectively. Seizure phenotypes are indicated according to the ILAE 2017 expanded classification. Neurological symptoms already present before seizure onset are highlighted in bold, neurological symptoms occurring during or after seizure onset are highlighted in italics. Effective anticonvulsants are highlighted in bold. GO = generalized onset. FO = focal onset. CBZ = carbamazepine; CLO = clobazame; GAB = gabapentine; LAC = lacosamide; LEV = levetiracetam; PHB = phenobarbitone; PIR = piracetame; PRED = prednisolone; TOP = topiramate; VIG = vigabatrine; VPA = sodium valproate. HIE = hypoxic-ischaemic encephalopathy. y = years; m = months; ND = no data

Corresponding EEG abnormalities typically featured three stages: i) initial multifocal sharp waves (Fig S4D), ii) short compound paroxysmal spike wave patterns (Fig 3D and Fig. S4E-G), iii) focal or generalized slowing with temporooccipital emphasis (Fig. S4E-G). In one case we could demonstrate how an initially normal EEG progressed to typical hypsarrhythmia (Fig. 3D).

Half of the patients showed an effective response to levetiracetam, the most commonly used anti-epileptic drug in pediatric epilepsy patients due to its well-established safety and efficacy profile. In cases where levetiracetam did not achieve seizure control, a range of other drugs including clobazam, gabapentin, valproate, carbamazepine, topiramate, and/or lacosamide proved also ineffective. Substantially improved seizure control was achieved with the ketogenic diet (KD) in patient III.1, commenced after an unsuccessful levetiracetam trial.

Symptoms suggestive of neurodegeneration including acquired dystonia, spasticity and behavioural changes preceded or occurred concomitantly with the onset of epilepsy, and remained relatively stable or, less frequently, progressed explosively, with more dramatic presentations such as opisthotonos often coinciding with bouts of systemic infection. This is of special relevance in *EPG5*-related VS, given the associated primary immunodeficiency [36].

## Discussion

Our study shows that *epg5* deficiency in *Drosophila* leads to a characteristic age-related neurodegenerative phenotype, recapitulating some of the aspects of VS, while neurodevelopmental defects, which are prominent in the majority of patients, are not evident. All the anomalies observed when modulating *epg5* in *Drosophila* are age-related, and develop in parallel to proteotoxic stress due to autophagic defects, indicating that the latter are likely to be the ultimate cause of most *epg5* neurological phenotypes in flies. The lack of robust neurodevelopmental defects in *epg5^ko^* flies in contrast to VS patients may be the result of the widely different developmental conditions between flies and humans. However, a mouse knock-out model for Epg5 [37] also failed to reproduce the full developmental impact of *epg5* mutations in VS patients, suggesting a physiological mechanism specific to human development as the key distinguishing factor.

The absence of neurodevelopmental defects in *epg5* mutants that nevertheless display clear seizure-like behaviours has led us to postulate that *EPG5*-related epilepsy in several patients may also be a consequence of neurodegenerative progression as a direct outcome of the proteotoxic stress due to autophagy impairment rather than of the primary neurodevelopmental defect. Indeed, several observations from our patient cohort support this hypothesis: In particular the onset of seizures from late infancy to childhood and the evolution of seizure types from focal-onset towards generalized-onset tonic or atonic seizures with loss of clonic components indicate a progressive course that is likely to be largely independent of the neurodevelopmental disorder. Moreover, the evolution of EEG abnormalities towards slow waves is detected at various stages in several common disease entities with a progressive neurodegenerative course, including Alzheimer’s dementia, Parkinson’s disease, or primary progressive aphasia [38–40]. Whilst we cannot exclude a contribution of the typical *EPG5*-associated neurodevelopmental brain MRI abnormalities, our findings strongly support a superimposed age-dependent neurodegenerative component, most likely associated with proteotoxic stress and the primary autophagy defect, linked to epileptogenesis not only in *Drosophila* but also in humans.

These results support the use of *Drosophila* as a model to study human epilepsies and a suitable replacement for vertebrate animals for basic studies with the potential to achieve substantial biomedical impact. In addition, we report here two novel methodologies, which allow automated higher-throughput and unbiased analysis of seizure-like behaviour and clear association of this behaviour with neuronal hyperactivity in live adult flies, strengthening this invertebrate model in the context of epilepsy research.

Our *Drosophila* work also supports the notion that caloric restriction mimetics could be considered in addition to more common pharmacologic treatments. A well-established form of dietary intervention in monogenic neurodegenerative diseases and epilepsies is the ketogenic diet (KD), which can induce autophagy in different settings [41–43] and is also protective in rat models of induced seizures [44]. Indeed, the only patient within our cohort who was commenced on a ketogenic diet after other anticonvulsants were ineffective showed a remarkable response not only in terms of epilepsy control but also in terms of other aspects of the disease phenotype. This suggest that autophagy stimulating therapies may be beneficial as a non-pharmacological alternative for seizure-control in patients with VS and related disorders.

There is clinical evidence for an *EPG5*-associated disease spectrum, ranging from classic VS to much milder neurodevelopmental disorders where epilepsy may be a prominent feature. While the response to fasting may depend on the extent of the autophagy defect in patients, this first report of a disease-modifying effect through dietary intervention in both *Drosophila* and humans suggest a potential therapeutic option for this currently incurable disease that surely deserves further clinical attention and investigations..

## Materials and Methods

### *Drosophila* strains and husbandry

Experiments were performed on *Drosophila melanogaster* (fruit flies) that were maintained in a 25°C incubator at a 12h light/dark cycle on standard fly food (0.8% w/v agar, 2% w/v cornmeal, 8% w/v glucose, 5% w/v Brewer’s yeast, 1.5% v/v ethanol, 0.22% v/v methyl-4-hydroxybenzoate and 0.38% v/v propionic acid). If the flies were calorically restricted, they were maintained on diluted standard fly food with a 1% agar solution in different volumes. Genetic crosses were made using virgin female flies and male flies and were kept at 25°C or at 18°C if the temperature sensitive Gal80 driver (Gal80ts) was used.

The following fly stocks were obtained from the Bloomington collection: w1118 (BDSC_3605), elavGal4 (BDSC_8765), tubGal80 (BDSC_5192), *UAS-snap25^IR^* (BDSC_27306), *UAS-CaLexA* (BDSC_66543) and *UAS-GFP^IR^* (BDSC_9331). Flies with a knockout of *epg5* and their respective controls were generated by BestGene Inc in a background line that was also obtained from Bloomington (BDSC_78781). From the VDRC collection *UAS-epg5^IR^* flies (VDRC_KK110420 (KK), VDRC_GD17469 (GD1), VDRC_GD17971 (GD2)) were used. *LexAop-luc* was a gift by Prof. Ralf Stanewsky (University of Münster, Germany) and Dr Alessio Vagnoni kindly gifted us *nSybGal4* flies. *UAS-GFP-Atg8a* was a gift by Tom Neufeld (University of Minnesota, USA). The *ubiGal80ts* flies were generated in the Fanto lab and previously described [45].

### CRISPR/Cas9 *epg5*^ko^ generation

The plasmid pCFD4 [46] was assembled through Gibson Assembly cloning using the following primers (gRNA sequences underlined):

Forward:

TATATAGGAAAGATATCCGGGTGAACTTCG<u>CGCTAAGATCGTGTACCGGC</u>GTTTTAGAGCTAGAAATAGCAAG;

Reverse:

ATTTTAACTTGCTATTTCTAGCTCTAAAAC<u>ACTCTTCTGACGGTATGCGA</u>CGACGTTAAATTGAAAATAGGTC

The pTV3 vector containing the *epg5* 5’ and 3’ recombination arms on both sides of the mCherry gene was assembled through PCR on genomic DNA and standard restriction cloning using the following primers

5’arm Forward: TCGCATGCAGGAGTAACGTCGCAAGAGC

5’arm Reverse: ATGCGGCCGCCGGGCGAGTTTAGGTCAAGT

3’arm Forward: GCACTAGTGCTGGATAGCATCGATGGGG

5’arm Reverse: GTACCGGTGTCCCAGACCTCTGGAATGG

PCR validation was obtained using the following primers

5’ *epg5* Forward: TTTAACATCCTGCGCTGTCCC

5’ *mCherry* Reverse: GAAGCGCATGAACTCCTTGAT

### Lifespan assay

Newly eclosed flies were collected daily for 3 days and then transferred to a 29°C incubator with humidity control in batches of 20 flies per vial (equal mix of males and females). Three times per week the flies were counted and flipped into a new vial with fresh fly food using CO_2_ to anaesthetise the flies. The number of flies that was still alive was recorded each time and flies that escaped or were stuck in the food were censored. A minimum of 60 flies per genotype was assessed.

### Negative geotaxis assay

Flies were collected and maintained in the same way as for the lifespan assay. In the morning of the assay 5 female flies per group were transferred to an empty tube that has markings indicating each centimetre starting from the bottom. The flies were allowed to recover from the anaesthesia for a minimum of 30 minutes before performing the assay. Using a custom-made setup, the tubes were placed in a holder which was gently banged down on a table to induce a negative geotaxis reaction from the flies. After 1 minute this was repeated, at a total of 7 times. All assays were videorecorded and analysed afterwards by measuring the distance the flies had climbed during the 20 seconds after the induction. The first and last repeat were excluded from the analysis and the average of the 5 flies per tube over the 5 repeats was recorded.

### DART assay

The Drosophila ARousal Tracking (DART) system was used to measure open field locomotor activity of the flies [28]. Flies were collected as described for the lifespan assay and anaesthetised using ice. They were individually placed in an arena of 35 mm diameter and 1.8 mm height on a platform. The flies were recorded by a camera for the total duration of the assay. After a resting period of 30 minutes the flies were mechanically stimulated through vibration of the platform every 15 minutes (5 vibrations, 0.2 seconds long, 0.5 seconds apart). This mechanical stimulation was repeated 5 times and the assay was finished with a final 15-minute rest period.

### Epi-DART prototype and analysis

The epilepsy thermo-hydraulic chamber and the analysis protocols have been developed in collaboration with BFKLab LTD and will be thoroughly described elsewhere. Here we report briefly the protocol used. Flies were anesthetised on ice and a single fly was inserted into small plastic trikinetics tube sealed with impermeable caps. The chamber’s volume is around 90ml and the pump hydraulic system has a capacity of 85ml/min which is manually controlled with a proportional– integral–derivative controller. These ensure acute thermal change. After 10 min recovery at RT (22°C) 10 tubes were inserted rapidly in the thermic chamber pre-heated at 41°C. The 41 °C heat shock was induced for 2 mins and flies’ behaviour was recorded. Flies during convulsions due to seizing activity drop at the bottom of the tube and remain there until recovery. Five flies per sex per genotype per batch and total 8 batches per sex were tested swapping side of the chambers. A seizure event was defined by the following parameters: duration >0.5 s; Position of the fly for the entire duration: 0-2.5 mm (bottom of the tube); Speed: >3 mm/s. Data collection and analysis was performed by the DART software.

### Hyperthermia-induced seizure assay

Flies were exposed to hyperthermia to measure seizure-like behaviour [25,29]. Flies were individually transferred into empty vials using CO_2_, followed by a recovery period of minimum 30 minutes. During the assay the vials were submerged in water with a temperature of 42 ± 0.5°C for 2 minutes. The seizure start time and recovery time of the flies was recorded. Seizure-like behaviour was defined as twitching of the legs, erratic movements, and the loss of their upright position, frequently followed by a period of paralysis. Recovery was defined as being able to maintain an upright position again and to walk around in the vial. All assays were performed blinded for the genotype.

### Luciferase assay

In the CaLexA system a modified NFAT (UAS-mLexA-VP16-NFAT, hereafter referred to as UAS-CaLexA) is used which is driven by the UAS-Gal4 system and is in this case expressed pan-neuronally by using the elavGal4 driver. Once imported in the nucleus NFAT binds to the lexAop promotor that drives transcription of the luciferase enzyme. This luciferase then oxidates the luciferin compound (D-luciferin, monosodium salt, Thermofisher), which is administered to flies orally. The flies were submitted to the hyperthermia-induced seizure assay and then transferred to a 96-well plate containing a luciferin-agar-sucrose solution (25 mM D-luciferin, 1% agar, 4% sugar). The plate was then transferred to a LB962 CentroPRP Microplate Luminometer (Berthold Technologies, UK) and photon emission was recorded for 24 hours. All assays were performed blinded for the genotype.

### Immunofluorescence

Adult flies were anaesthetised on ice and brains were dissected in PBS-T (phosphate-buffered saline with 0.3% Triton X-100). The brains were fixed in 4% paraformaldehyde for 30 minutes in PBS, followed by a 30-minute wash in PBS-T and blocked in 5% normal goat serum in PBS-T for 30 minutes. They were then incubated with the primary antibody at 4°C overnight. The next day the brains were washed 3 times with PBS-T for 10 minutes at room temperature and afterwards incubated with the secondary antibody overnight at 4°C. Lastly, the brains were washed 3 times in PBS-T for 10 minutes, followed by incubation in Hoechst (1 μg/ml) for 10 minutes, a wash with PBS-T again and then they were mounted on a slide using DAKO mounting medium. All images were taken using the A1R confocal microscope (Nikon). Antibodies used: Rabbit α-Ref(2)P (1:1000, kind gift by Tor Erik Rusten); Rabbit a-Atg8a (1:500, Merck ABC974), mouse anti-Fasciclin II (1D4 Developmental Studies Hybridoma Bank, 1:100); rabbit anti-Tyrosine Hydroxylase (ab128249 Abcam, 1:500). Secondary goat anti-mouse and anti-rabbit AlexaFluor 488 or 647 (Life Technologies, 1:1000).

### Quantification and statistical analysis

All data were analysed using Microsoft Excel and GraphPad Prism 9. Lifespan and seizure prevalence were analysed with the log-rank (Mantel-Cox) test. Unpaired Student’s t-test was used for comparison of 2 groups of normally distributed data. If the data were not normally distributed a Mann-Whitney test was performed instead. Two-way ANOVA with Bonferroni’s or Dunnett’s multiple comparisons test was used for comparison of 2 groups of normally distributed data confounded by a third parameter (e.g., different timepoints).

### Deep phenotyping and genomic investigations

Patients were recruited from multi-center studies via established collaborations, the web platform GeneMatcher [47], and a national study on rare pediatric neurological diseases in Germany (ESNEK). All parents gave informed consent. The study was approved by the ethics committee of the Medical Faculty, University of Cologne (20-1711) and other regional institutional review boards [48]. Massively parallel sequencing was performed as previously published [49]. Dideoxy sequencing was performed to test for co-segregation of variants with the disorder. Deep phenotyping of clinical features, patient and family histories, electroencephalography (EEG) recordings and brain magnetic resonance imaging (MRI) studies were carefully conducted and reviewed by a group of pediatric neurologists, clinical geneticists, and neuroradiologists [48].

## Acknowledgements

We thank BFKLab for its assistance with the behavioural assays, the analysis and the continuous support. We thank Alessio Vagnoni, the VDRC and BDSC for fly stocks. We thank Best Gene for fly embryo injection. We thank Jean Paul Vincent and Cyrille Alexandre for the pTV3 and pCFD4 plasmids. We thank Tor Erik Rusten for the Ref(2)P antibody. This work was supported by grants from the European Union’s Horizon 2020 Research and Innovation Programme (765912 — DRIVE — H2020-MSCA-ITN-2017) to CD and HJ, Action Medical Research (2446) to HJ and MF, the National Centre for the Replacement, Refinement & Reduction of Animals in Research (NC3Rs) (NC/V001051/1) to MF and JJ. HSD was supported by the Cologne Clinician Scientist Program / Medical Faculty / University of Cologne and German Research Foundation (CCSP, DFG project no. 413543196). The authors declare that they have no conflict of interest.

