## Supplemental Figures for "Epg5 links proteotoxic stress due to defective autophagic clearance and epileptogenesis in *Drosophila* and Vici Syndrome patients"

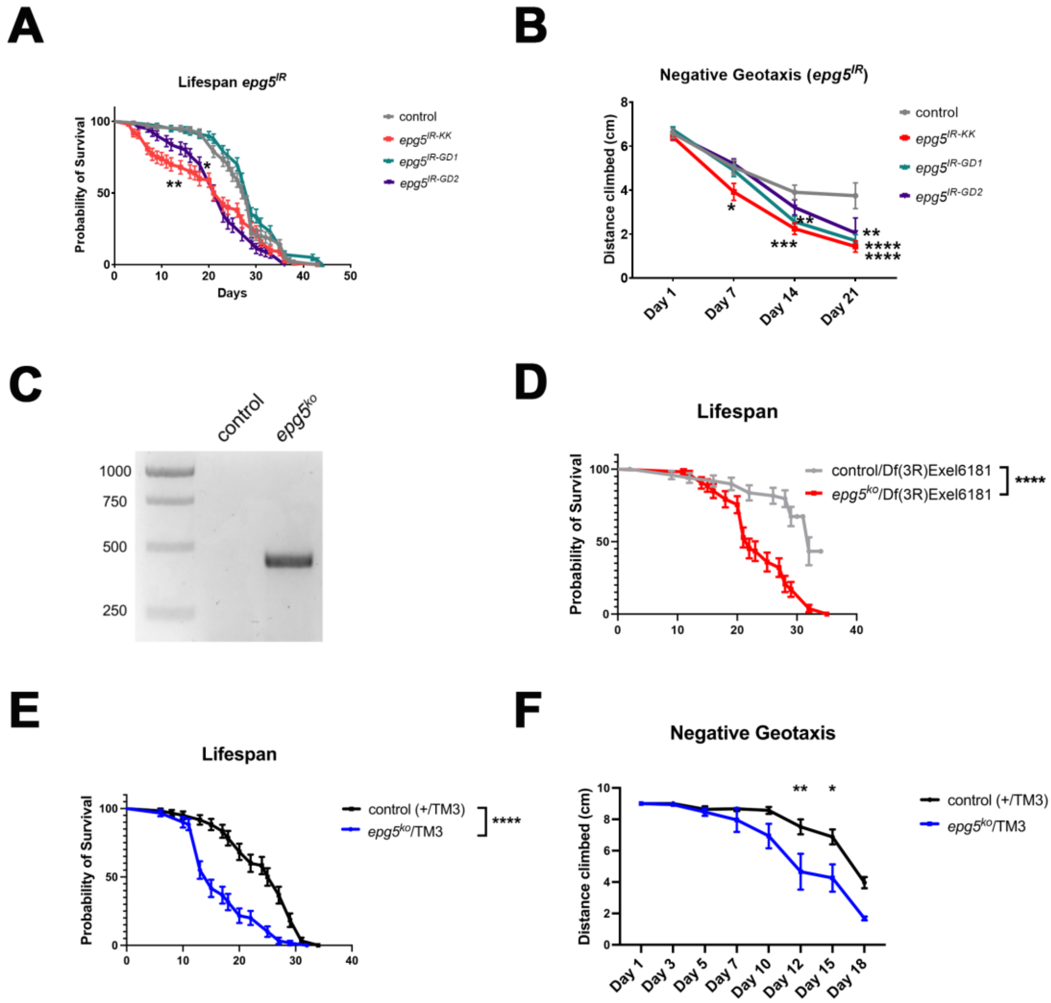


### Deneubourg *et al.* Figure S1

**Figure S1. Age-related degeneration in *epg5* knock down and knock out flies.** (A) Survival rate (%) of flies expressing three different *UAS-epg5*$^{IR}$ transgenes in the nervous system with the *elav-Gal4* driver in comparison to *UAS-GFP*$^{IR}$ control. N=60 flies per genotype. Kaplan Meier Log-rank test, *p<0.05, **p<0.01. (B) Climbing assay of flies of the same genotype as in A. N=~30 flies per genotype. Data are presented as mean ± S.E.M, Two-way ANOVA with Dunnett's multiple comparisons test. *$p<0.05$, **$p<0.01$, ***$p<0.001$, ****$p<0.0001$. (C) PCR on the 5' arm of the recombinant gene with a primer matching the *epg5* gene and one primer matching the *mCherry* sequence. (D) Survival rate (%) of *epg5*$^{ko}$ flies in comparison to an isogenic control, both trans-heterozygous over Df(3R)Exel6181, a deficiency removing the entire *epg5* locus. N=50-53 flies per genotype. Kaplan Meier Log-rank test, ****$p<0.0001$. (E) Survival rate (%) of *epg5*$^{ko}$ heterozygous flies in comparison to an isogenic control both carrying a TM3 chromosome balancer. N=60 flies per genotype. Kaplan Meier Log-rank test, ****$p<0.0001$. (F) Climbing assay of *epg5*$^{ko}$ heterozygous flies in comparison to an isogenic control both carrying a TM3 chromosome balancer. N=~30 flies per genotype. Data are presented as mean ± S.E.M, Two-way ANOVA with Bonferroni's multiple comparisons test. *$p<0.05$, **$p<0.01$.

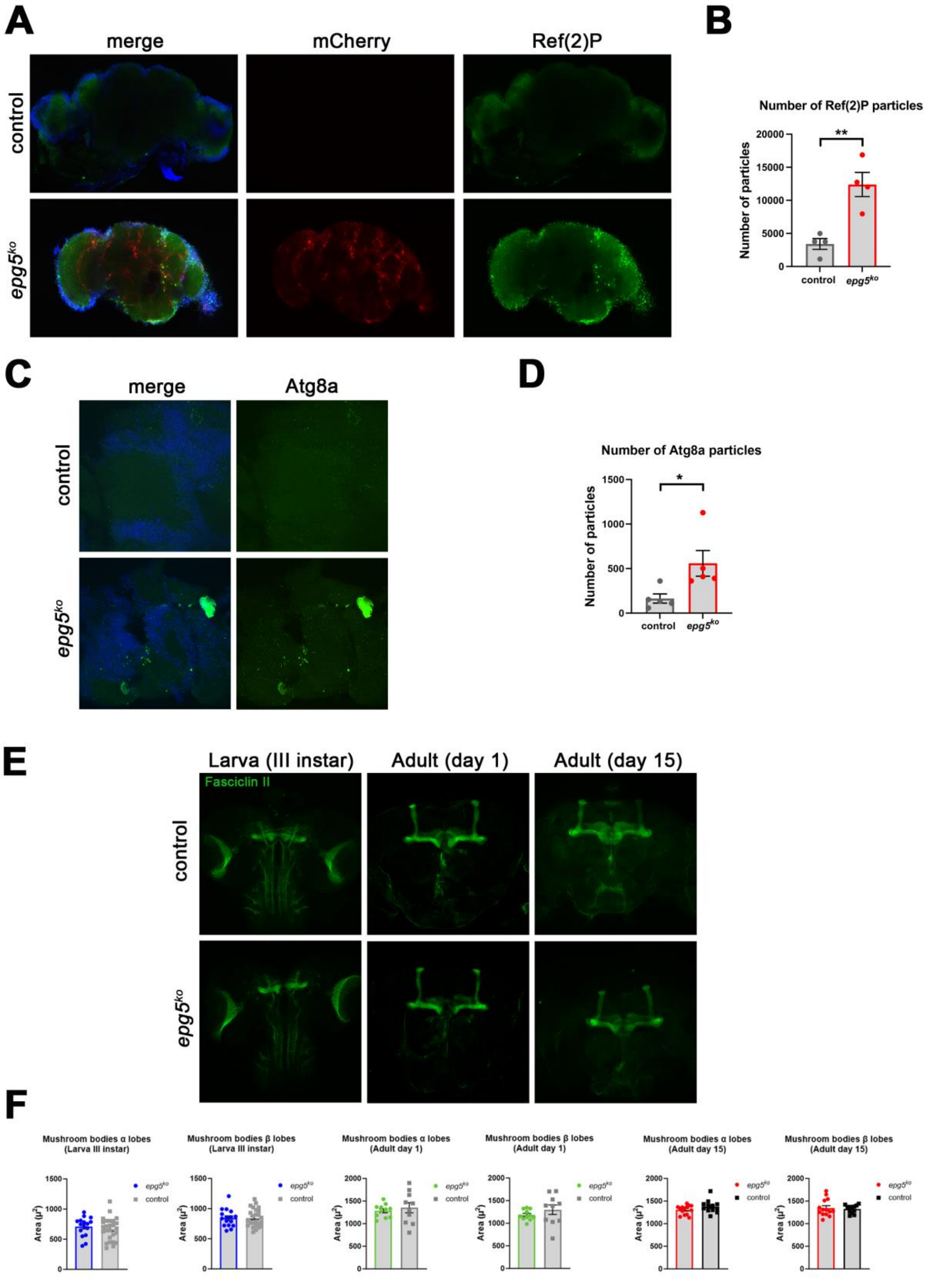


### Deneubourg *et al.* Figure S2

**Figure S2. Autophagy and neurodegeneration in *epg5* knock out flies.** (A) Representative images of dissected brains from $epg5^{ko}$ and isogenic control flies. mCherry fluorescence in red. Ref(2)P in green. A DAPI staining was used to show the outlines of the brains. (B) Quantification of the average number of Ref(2)P particles per brain. N=4 brains per genotype. Data were analysed using an unpaired

Student's t-test. **p<0.01. (C) Representative images of dissected central brains parts from *epg5*$^{ko}$ and control flies and stained for Atg8a (green) and DAPI (blue). All flies were expressing a *UAS-GFP-Atg8a* transgenes in all neurons with the *elavGal4* driver. (D) Quantification of the average number of Atg8a puncta per brain. N=5 brains per genotype. Data were analysed using an unpaired Student's t-test. *p<0.05. (E) Max intensity z-projections of confocal stacks through the brains of *epg5*$^{ko}$ and isogenic controls at different life stages stained for Fasciclin II (green) to reveal mushroom body projections. (F) Quantification of the area covered by the mushroom body projections in $\alpha$ and $\beta$ lobes in *epg5*$^{ko}$ and isogenic controls at different life stages. No difference is observed at any stage in these neurons.

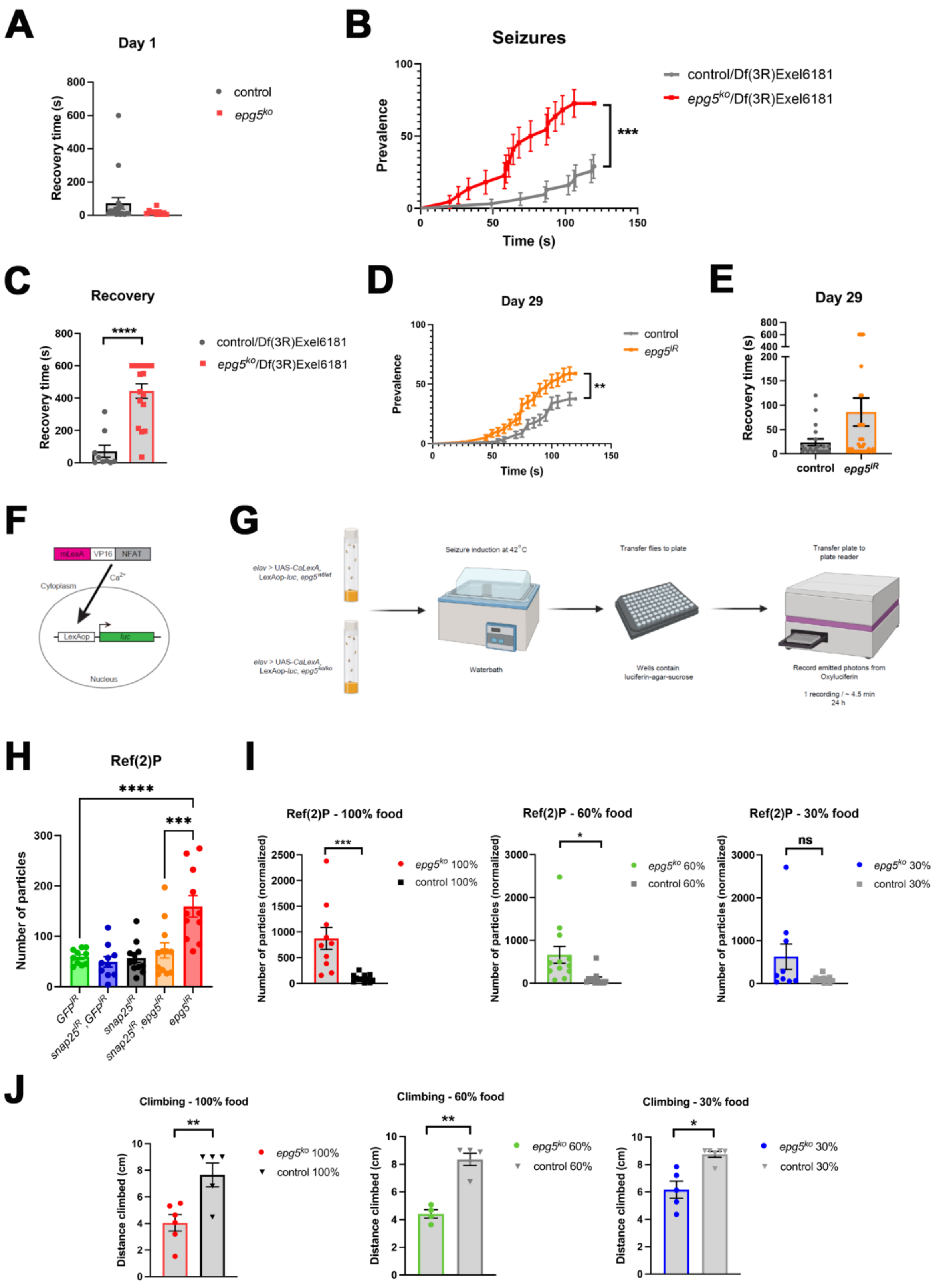


### Deneubourg *et al.* Figure S3

**Figure S3. Seizure-like behaviour in epg5 mutant and knock down flies.** (A) Recovery time of *epg5*$^{ko}$ flies in comparison to an isogenic control with seizures at day 1. N=10-18 flies per genotype. Data are presented as mean ± S.E.M. (B) Seizure prevalence in *epg5*$^{ko}$ flies in comparison to an isogenic control, both trans-heterozygous over Df(3R)Exel6181, a deficiency removing the entire *epg5* locus. N=22-31 flies per genotype. Data are presented as mean ± S.E.M and analysed using the Log-rank (Mantel-Cox) test, ***p<0.001. (C) Recovery time of flies with seizures in B. N=9-17 flies per genotype. Data are presented as mean ± S.E.M and analysed using an unpaired Student's t-test ****p<0.0001. (D) Seizure prevalence of *epg5* knock down with the KK line expressed though *nSyb-Gal* driver in the nervous system and related control. Data are presented as mean ± S.E.M. and are analysed using the Log-rank (Mantel-Cox) test. N=18-24 per genotype. **p<0.01. (E) Recovery time of *epg5* knock down with the KK line expressed though *nSyb-Gal* driver in the nervous system and related control. N=20-30 flies per group. Data are presented as mean ± S.E.M. (F) Schematic Diagram of the CaLexA-Luciferase method. (G) Schematic representation of the CaLeA-Luciferease protocol. (H) Quantification of the average number of Ref(2)P particles per brain from *snap25* and or *epg5* knock down in the nervous system and related controls. Brains were stained with an anti-Ref(2)P antibody and DAPI, to detect the brain outlines. Data are presented as mean ± S.E.M. and analysed through a One-Way ANOVA with Tukey's post-hoc test for multiple comparisons. N=10-12 flies per group. ***p<0.001, ****p<0.0001. (I) Quantification of the average number of Ref(2)P particles per brain in $epg5^{ko}$ and isogenic control flies maintained on a standard diet (100% food), calorically restricted with 60% food or with 30% food. Brains were stained with an anti-Ref(2)P antibody and DAPI, to detect the brain outlines. Data are presented as mean ± S.E.M. and are analysed using an unpaired Student's t-test. N=9-12 flies per group. ***p<0.001, *p<0.05. (J) Climbing assay of $epg5^{ko}$ and isogenic control flies maintained on a standard diet (100% food), calorically restricted with 60% food or with 30% food. Data are presented as mean ± S.E.M. and are analysed using an unpaired Student's t-test. N=20-30 flies per group. *p<0.05, **p<0.01.

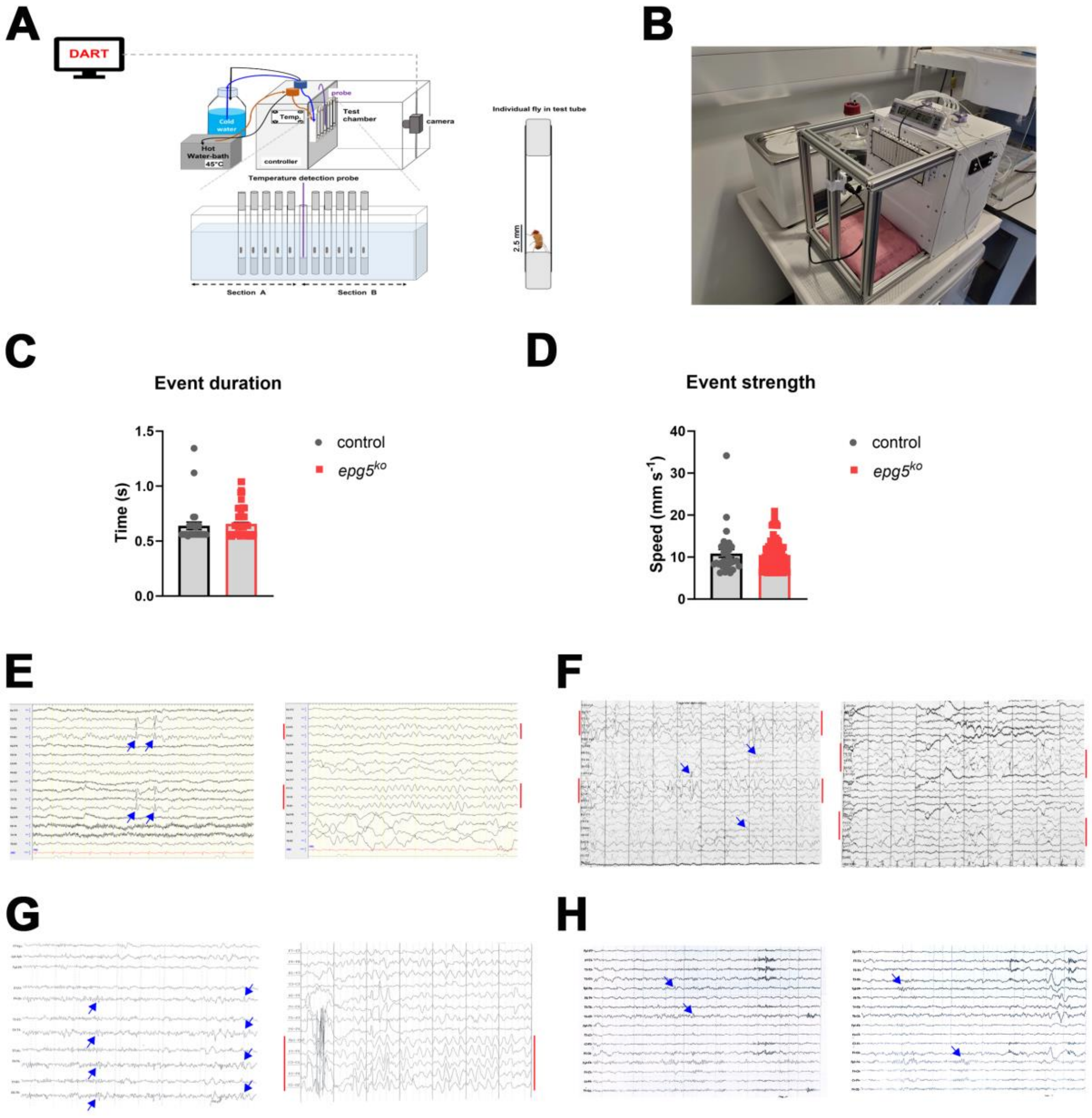


### Deneubourg *et al.* Figure S4

**Figure S4. Epi-DART and EEG analysis of seizures in Drosophila and in patients with *EPG5*-related epilepsy** (A) Schematic representation of the epi-DART apparatus (B) picture of the epi-DART thermohydraulic chamber prototype. (C) Average seizure event duration as calculated by the DART software. N=28-80. (D) Average seizure event strength as calculated by the DART software. N=28-80.

(E) EEG recordings from Patient I.2. with focal aware motor onset tonic seizures with paresis showing spike wave complexes (blue arrows) at age of two years (left panel), and progression to slow waves (channels marked red) at age of four years (right panel). (F) EEG recording from Patient IV.1 with generalized onset motor atonic seizures showing high voltage sleep recording and independently right-sided slow bursts (channels marked red) (left panel), and bilateral semirhythmic spikes and sharp waves with maximum left posterior parasagittal (blue arrows, right panel) and slow waves (channels marked red) at age of one year and 11 months. (G) EEG recordings from Patient V.1 with focal onset motor tonic seizures showing only mild abnormalities at age of three years (left panel, blue arrows indicate two sharp waves and spike waves), and progression to generalized slow waves (right panel, channels marked red) pronounced over the left hemisphere following development of generalized onset motor spasms at age five years. (H) EEG recording from Patient VI.I with focal onset motor clonic seizures showing multifocal sharp waves (blue arrows) at age of four years.
